# Burden-driven feedback control of gene expression

**DOI:** 10.1101/177030

**Authors:** F Ceroni, S Furini, TE Gorochowski, A Boo, O Borkowski, YN Ladak, AR Awan, C Gilbert, GB Stan, T Ellis

## Abstract

Cells use feedback regulation to ensure robust growth despite fluctuating demands on resources and different environmental conditions. Yet the expression of foreign proteins from engineered constructs is an unnatural burden on resources that cells are not adapted for. Here we combined multiplex RNAseq with an *in vivo* assay to reveal the major transcriptional changes in two *E. coli* strains when a set of inducible synthetic constructs are expressed. We identified that native promoters related to the heat-shock response activate expression rapidly in response to synthetic expression, regardless of the construct. Using these promoters, we built a CRISPR/dCas9-based feedback regulation system that automatically adjusts synthetic construct expression in response to burden. Cells equipped with this general-use controller maintain capacity for native gene expression to ensure robust growth and as such outperform unregulated cells at protein yields in batch production. This engineered feedback is the first example of a universal, burden-based biomolecular control system and is modular, tuneable and portable.

## INTRODUCTION

Maintenance and expression of synthetic constructs adds an unnatural load to host cells that is typically known as burden ^1^. While some burden can stem from the specific roles of the genes in the constructs (e.g. enzymes that consume metabolites), most burden placed on cells is simply due to the consumption of finite cellular resources, such as polymerases and ribosomes during expression of the construct genes ^2 3 4^. The cellular response to the load of extra expression is typically decreased growth and global physiological changes. These are somewhat unpredictable and typically reduce the expected performance of cells hosting synthetic constructs ^5 6 7 8^.

Efforts to increase control and predictability of engineered biological systems has led to tools to measure and reduce burden which have come about through improvements in our understanding of host-construct interactions ^9 10^. This has led, among other things, to the development of new orthogonal expression systems ^11^, a transcriptional resource allocator able to tune the transcriptional capacity available for a synthetic system ^12^, and libraries of promoters able to tune the expression of burdensome proteins and decrease cellular stress 13. These are all strategies that bypass some of the resource-sharing required for gene expression but do not fully-eliminate it.

In parallel, mathematical modelling has enabled researchers to understand the main bottlenecks involved in expression burden and consider methods to alleviate these ^14 15 4^. Previously, we developed a *capacity monitor* system for *E. coli* that quantifies the capacity for gene expression taken from the cell by synthetic constructs. This was used to identify more efficient construct design strategies that showed robust growth with less chance of mutations arising in populations ^16^. In that work, we noted that a large competition for expression resources, especially free ribosomes, causes decreased growth rate of the host *E. coli* cells over the next few hours. As sudden overexpression of extra genes is likely to be encountered naturally in bacteria (e.g. upon plasmid uptake or phage infection), we reasoned that cells must have native mechanisms to adapt and reallocate resources to continue growth. Investigating how *E. coli* adapts in the face of synthetic construct expression offers the chance to identify native systems that sense and respond to burden and exploit these downstream in synthetic gene circuits.

To identify a host cell response to burden requires looking at all gene expression before and after burden is triggered. RNA sequencing (RNAseq) allows semi-quantification of all the transcription within cells at a given moment in time ^17 18^ and differential RNAseq (comparing sequencing read numbers between conditions) has been used to characterise transcriptional changes in response to environmental and growth conditions, enabling identification of differentially-expressed genes with high accuracy ^19 20^. Here, we paired our previously described *in vivo* capacity monitor assay for measuring expression burden in *E. coli* with a multiplex RNAseq strategy in order to explore the transcriptional changes that occur in the host after the onset of burden from expression of induced synthetic constructs. This enabled us to gain an improved understanding of the burden of a set of exemplar synthetic constructs and to identify common regulated promoters that quickly activate or repress transcription due to the burden of extra gene expression. Using one of these promoters we then built a tunable CRISPR/dCas9-based feedback system that can regulate synthetic construct expression in response to burden, thereby maintaining cellular capacity during synthetic expression. This system is the first example of a general burden-based biomolecular feedback control system that provides increased robust performance in terms of protein yield from synthetic constructs and host cell growth.

## RESULTS

In this study, we first investigated the *in vivo* burden and transcriptome response of *E. coli* strains expressing a chosen set of exemplar, inducible synthetic constructs with different genes, codon-optimisation and regulation (**Figure 1a**). Two strains, DH10B and MG1655, were chosen as hosts, with both incorporating the capacity monitor described previously ^16^. In these cells, a small expression cassette is stably integrated into the chromosome to produce GFP at a constitutive rate. Inducing heterologous gene expression, such as from plasmids, burdens the cell by depleting the host’s resources and this can be measured from the subsequent decrease in the GFP production rate per cell.

For the synthetic constructs, we focused on 3 different exemplar cases of plasmid-based inducible expression; (i) inducible reporter expression, (ii) overexpression of a large non-functional heterologous protein, and (iii) expression of an operon encoding a metabolic pathway. For the latter, we expressed the Lux operon from *Vibrio fischeri* (codon-optimised for *E. coli*) from the araBAD induction cassette (AraC-pBAD) on a high copy pSB1C3 plasmid (**pSB1C3-Lux**). The same plasmid backbone and a version of the same induction cassette, but with a mutation for slightly higher expression, were also used for expression of the large heterologous protein (VioB with C-terminal mCherry fusion) in a construct called **pSB1C3-H3** used in our previous study ^16^. The metabolite-induced reporter construct (**pD864-LacZ**) differed from the other two by being on a different plasmid backbone (pD864), having a different antibiotic selection (ampicillin vs chloramphenicol) and a different induction (rhamnose vs arabinose). It also only contained genes and promoter sequences native to *E. coli* (the pRhaBAD promoter and LacZ). With these three constructs, we also included empty plasmid constructs (pSB1C3 and pD864) as controls, and for preliminary investigations we also added a fourth synthetic construct called **pLys-M1**. This is an alternative design to pSB1C3-H3 that imposes more burden despite being on a lower-copy plasmid (pLys), due to having a stronger ribosome binding site (RBS) ^16^. In all cases externally-inducible gene expression was required to enable control of the start of burden, so that the very first transcriptional response and outcome can be tracked.

The four constructs and plasmid-only controls (**Figure 1b, Table S1**) were first characterised in the two different host strains of *E. coli* using capacity monitor measurement as an *in vivo* plate-based burden assay ^16^. By comparing GFP production rate per cell 1 hour after induction (or at the same time-point in samples with no induction) the burden of the constructs can be inferred (**Figure 1c**). A significant decrease in GFP production rate was observed for all samples with induced synthetic constructs, confirming that they all cause burden due to use of shared gene expression resources in the cells. Construct pLys-M1 clearly displays the greatest burden, whereas the least burden is seen with just the pD864 plasmid alone, which shows the same capacity as the untransformed host strains. The assay can also determine the growth rates of the cells during the experiment, and these were decreased after 1 hour of construct induction (**Figure S1**).

Having verified that the three types of construct all impose measurable burden, we next turned to multiplex RNAseq analysis for characterisation of the transcriptional performance of the constructs and their host cells. Growth of cells for RNA harvesting was done in a similar manner as for the *in vivo* assay but RNA samples were taken at 15 and 60 min post-induction in order to investigate rapid changes to the transcriptome. Total RNA harvested per sample closely correlated with the concentration of cells previously measured in the *in vivo* assay at equivalent time-points, giving us confidence that the RNAseq and *in vivo* assay datasets can be directly compared (**Figure S2**). To avoid confusing the cell’s response to burden with its response to the applied inducers (arabinose and rhamnose), the control samples used were the strains with empty plasmids, also exposed to the inducers for the same amount of time.

A total of 90 samples were multiplexed for the RNAseq (**Table S2**) following individual RNA purification, rRNA removal and other sample prep steps (see **Methods**). This number allowed three biological replicates for the constructs and controls, at the two time points and in two strains (DH10B and MG1655). Library preparation used a custom protocol adapted from previous Nextera kit methods ^21^. Total RNA was first extracted and assessed (RIN > 9), mRNA was enriched by rRNA removal ^22^, and retro-transcribed to cDNA. Nextera XT protocol was used for library synthesis utilising tagmentation to fragment cDNAs and attach adapter sequences. Limited-cycle PCR targeting the adapters amplified insert DNA and added barcodes for dual-index sequencing of pooled samples. All 90 samples were pooled and sequenced using two lanes of HiSeq 2500 sequencer with 100 bp paired-end reads.

**Figure 1.**
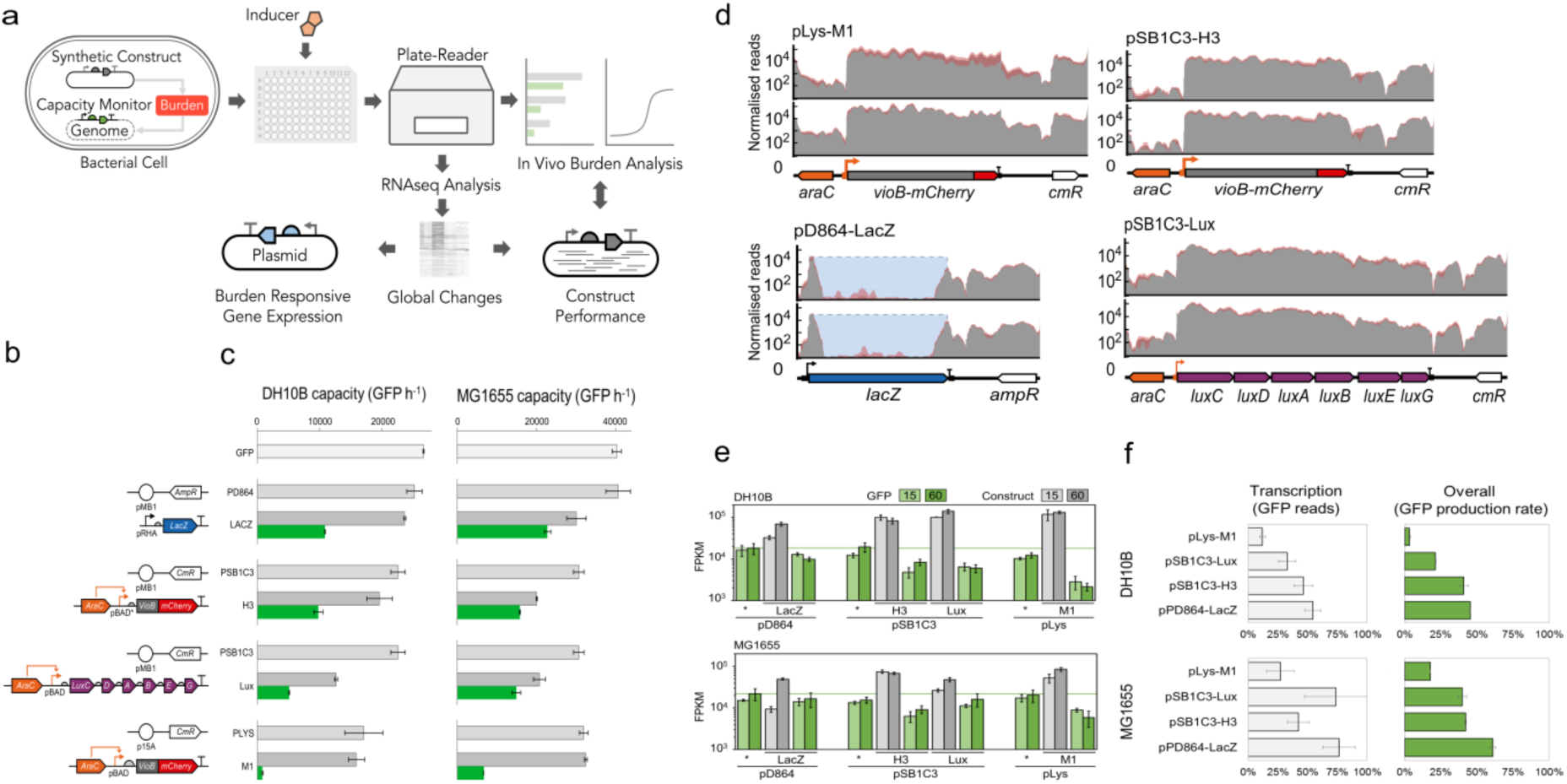
Characterisation of host-construct interactions in *E. coli*. **(a)** Workflow scheme of the present study. Cells transformed with different synthetic constructs and their empty plasmid control are used to measure the *in vivo* burden response and global transcriptional changes with RNAseq. Burden responsive promoters are identified and used in the design of burden-based biomolecular feedback systems. **(b)** Diagrams of the four constructs used in this study and their plasmid controls. pBAD and pRHA are inducible by arabinose and rhamnose, respectively. **(c)** *In vivo* cell capacity analysis 1 hour post-induction. Grey and green bars represent uninduced and induced samples, respectively. Darker grey bars represent the uninduced samples; lighter grey bars represent the empty plasmids. The lighter top bars represent the GFP production rate per cell in the untransformed strains. Error bars show standard deviation of two full independent repeats. **(d)** Transcription profiles of the synthetic constructs in DH10B cells 15 (top) and 60 (bottom) min post-induction. Traces show the normalised reads mapping to the plasmid constructs. Solid grey regions show average, and transparent red regions span from minimum to maximum of transcription profiles generated from three biological replicates. Genetic designs below each profile are drawn using SBOLv notation and show the precise location of each genetic part. For the pD864-LacZ construct, only reads uniquely mapping to the *lacZ* gene in the construct, and not the chromosomal copy, are used. Blue shaded regions denote the estimated expression level for the *lacZ* gene in the construct. **(e)** Transcriptional induction of the four constructs represented by mean FPKM values. Green bars are data for the calculated GFP FPKM and colour intensity relates to the time point; Grey bars are data for the FPKM corresponding to the construct gene; * denotes empty plasmid controls. Error bars show standard deviation of three independent repeats. **(f)** Burden imposed by the four constructs as measured by the percentage of available capacity. In the left panels, plots show GFP mRNA FPKM in construct samples compared to the pD864 control. In the right panels, plots show the same comparison but using GFP fluorescence data from the plate-based burden assay (Figure 1c). Error bars show standard deviation of three biological replicates.

Raw reads for all sequenced samples were quality assessed and trimmed. After assessment for potential batch effects, technical replicates were pooled. A FASTA format sequence file corresponding to the composite of strain, plasmid and integrated GFP capacity monitor cassette was constructed for each sample and used as reference for read alignment. All reads identified as unremoved rRNA were discarded and in the one case where reads could align to either the plasmid or the strain genome (for LacZ in pD864-LacZ in MG1655) the raw reads were assigned appropriately to match those of flanking sequence. Biological replicates were checked before generating the raw counts and this identified one outlier (first replicate of a DH10B empty strain control) which was discarded. Normalised FPKM counts were then generated and used for downstream analysis.

Normalised RNAseq data were first used to generate the transcriptional profiles of the plasmid constructs at 15 and 60 min post-induction (**Figure 1d**). The mapped reads that align to the plasmid sequences show the transcribed regions with the magnitude of their transcription revealing strong expression from the induced promoters, as well as expression from the selection markers and some pervasive transcription from other regions of the plasmids. Total plasmid-based transcription in the construct samples accounted for a significant percentage of all non-rRNA transcription in the *E. coli* (**Table S3, Table S4**). Indeed, only 15 min post-induction in DH10B cells, between 11.7% (pD864-LacZ) and 47.3% (pLys-M1) of all mapped reads came from the synthetic constructs. Notably, total transcription from the empty pLys plasmid was unexpectedly higher than for the other two empty plasmids (pD864 and pSB1C3) in both strains, and especially so in DH10B cells.

The RNAseq FPKM reads were next used as indicators of the transcriptional levels of both the GFP capacity monitor cassette and the output expression cassettes of the synthetic construct plasmids. At both 15 and 60 min post-induction, high levels of transcription from the constructs corresponds to a decrease in chromosomal GFP transcription when compared to values measured for the cells containing only control plasmids (**Figure 1e**). This demonstrates that induced transcription of synthetic constructs has a rapid effect on genomic transcription for all constructs tested and that burden clearly affects the host transcriptome. Indeed, when we directly compare the decrease in GFP RNAseq reads per sample to the decrease in GFP production rate per cell (as measured by the *in vivo* assay) after 1 hour we see very similar profiles (**Figure 1f**). This correlation between GFP production rate per cell and GFP FPKM reads per sample holds for most of the samples tested including controls (**Figure S3**).

Notably, the decrease in GFP transcription is least pronounced for pD864-LacZ, which expresses its output transcript (*lacZ* mRNA) significantly less than the other constructs express theirs. This is likely due to the activator of the pRhaBAD promoter, RhaS, only being provided for this construct *in trans* by the strain genomic copy, whereas the activator of the pBAD promoter, AraC, is provided in higher copy *in cis* as part of the plasmids for the other synthetic constructs. As with the *in vivo* assay of expression capacity, pLys-M1 clearly had the largest impact on host cell GFP transcription, and this was especially pronounced in DH10B cells. These naturally grow slower than MG1655 cells, and have lower gene expression capacity as measured by our assay. The DH10B strain also has a non-functional stringent response system ^23^.

Together the *in vivo* assay and RNAseq data reveal variation in the degree of burden imparted by the different constructs and some differences in behaviour in the two strains. Next, we investigated the effect that construct expression has on the rest of the cell’s transcriptome and specifically examined whether the cells had a common response to burden regardless of the genetic content of the synthetic construct or which strain was used. For every genomic gene except the ribosomal RNA genes, we calculated the fold change in gene expression between the empty plasmid sample and the construct, and from this we then determined the genes showing significant differential expression. The number of genes showing significant up- and down-regulation after 60 min is shown for the three main constructs pSB1C3-H3, pSB1C3-Lux and pD864-LacZ in both host strains (**Figure 2a**), with the same analysis also done for samples at 15 min (**Figure S4**). For the sake of clarity, the effects of pLys-M1 are excluded from these figures as this construct is simply a more burdensome version of pSB1C3-H3. However, equivalent analyses that include pLys-M1 are shown in **Figure S5** and **Figure S6**.

**Figure 2.**
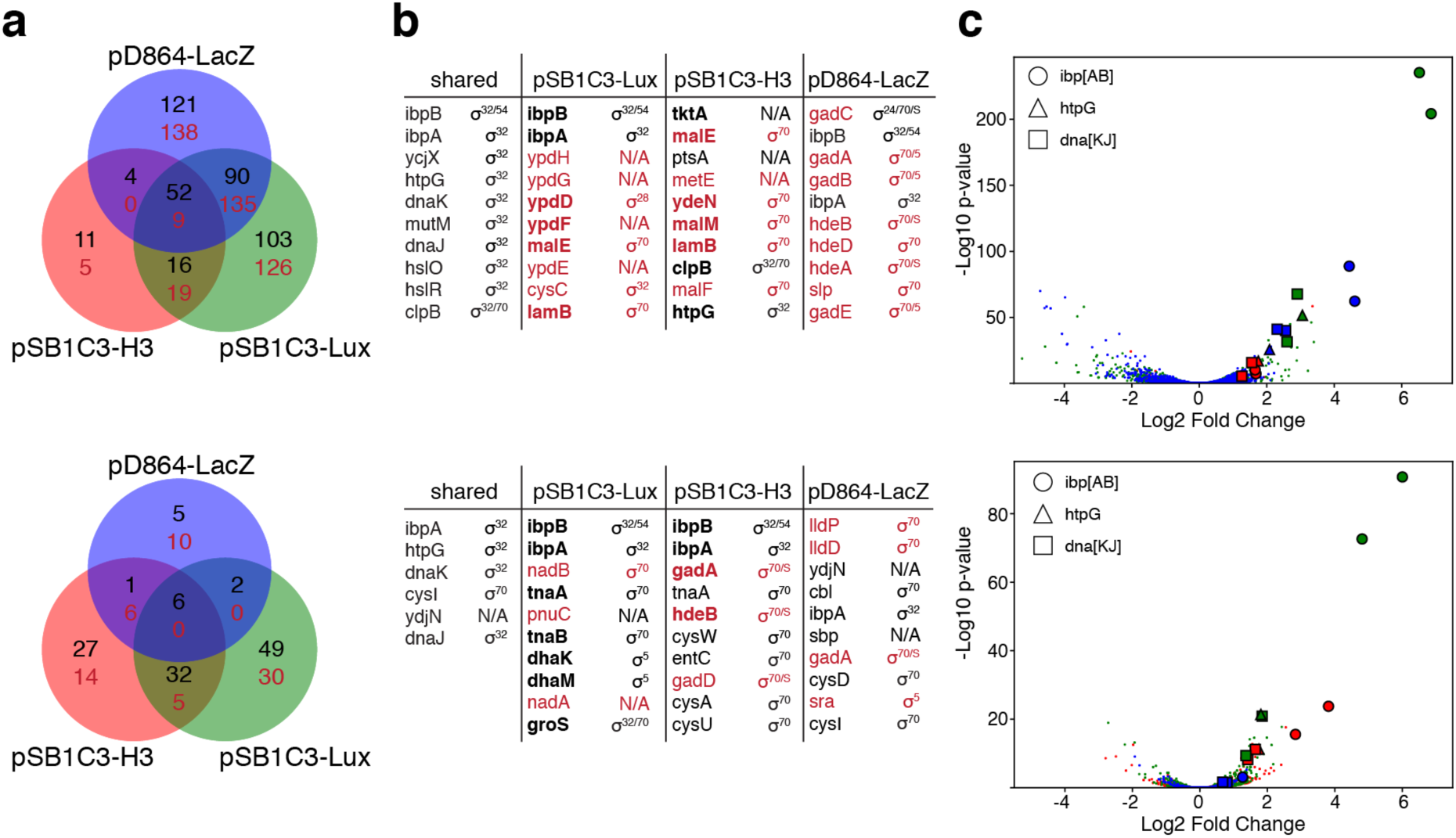
Global transcriptional changes in response to burden 60 min post-induction of synthetic construct expression. The set of significantly differentially expressed native genes (alpha < 0.05) was identified in DH10B (top panels) and MG1655 (bottom panels) cells separately for each of three synthetic gene circuits pSB1C3-Lux, pSB1C3-H3 and pD864-LacZ, comparing cells transformed with the synthetic circuits to cells transformed with the corresponding empty plasmids. **(a)** Venn diagrams of differentially expressed genes. Venn diagrams report the number of up-regulated and down-regulated genes for each sample in DH10B and MG1655, which are displayed in black and red, respectively; **(b)** List of top ten differentially expressed genes (black are up-regulated, red are down-regulated). Tables report the top ten differentially expressed genes for each construct and the associated regulatory sigma factors. Bold names indicate genes differentially expressed also at 15 min post-induction. Left columns show the differentially expressed genes that are shared among all 3 constructs; **(c)** Volcano plots. Volcano plots show genes that display large changes and are also statistically significant. Symbol colours correspond to the constructs colours displayed in (a) with pSB1C3-Lux, pD864-LacZ and pSB1C3-H3 displayed in green, blue and red, respectively.

Overall, the number of genes with statistically significant differential expression 60 min post-induction was higher in DH10B cells than in MG1655. While many of the genes were up-or down-regulated only with one construct, a significant number of these genes changed expression for two of the three constructs, and 61 genes in DH10B and 6 genes in MG1655 were differentially expressed in all three cases. The ten genes showing the greatest significant difference in expression for each construct were then examined to determine their regulation, by matching them to the sigma factor family known to direct their expression (**Figure 2b**). An equivalent list was also made for the top ten differentially expressed genes shared among all three constructs for both strains (**Figure 2b**). All these common genes were up-regulated in the samples with the induced constructs and nearly all expressed from promoters that require the heat-shock response sigma factor (σ32). Three promoters in particular showed up-regulation with all constructs in both strains: the *ibpAB* and *dnaKJ* operon promoters and the *htpG* promoter. When all differentially expressed genes were visualised on Volcano plots, the transcripts from these three promoters clearly displayed the most significant changes (**Figure 2c**).

The transcription of *dnaKJ, ibpAB* and *htpG* was next inspected at the genomic level, along with the *groSL* operon and the capacity monitor GFP cassette (**Figure 3a**). Transcriptional profiles clearly show that transcription of these four σ32-regulated genes increases by at least an order of magnitude following construct expression 60 min post-induction, while GFP transcription stays relatively high. By using the mapped-read numbers upstream and downstream of the promoters for these genes, we then determined the difference in transcription from these promoters in response to the burden of construct expression (**Figure 3b**). This showed increases in promoter activity ranging from 10- to 1000-fold in DH10B cells and from 3- to 300-fold in MG1655. These four promoters can therefore be characterised as intrinsic biosensors for synthetic construct-induced burden in *E. coli*, although the precise mechanisms that lead to their activation are not fully understood.

**Figure 3.**
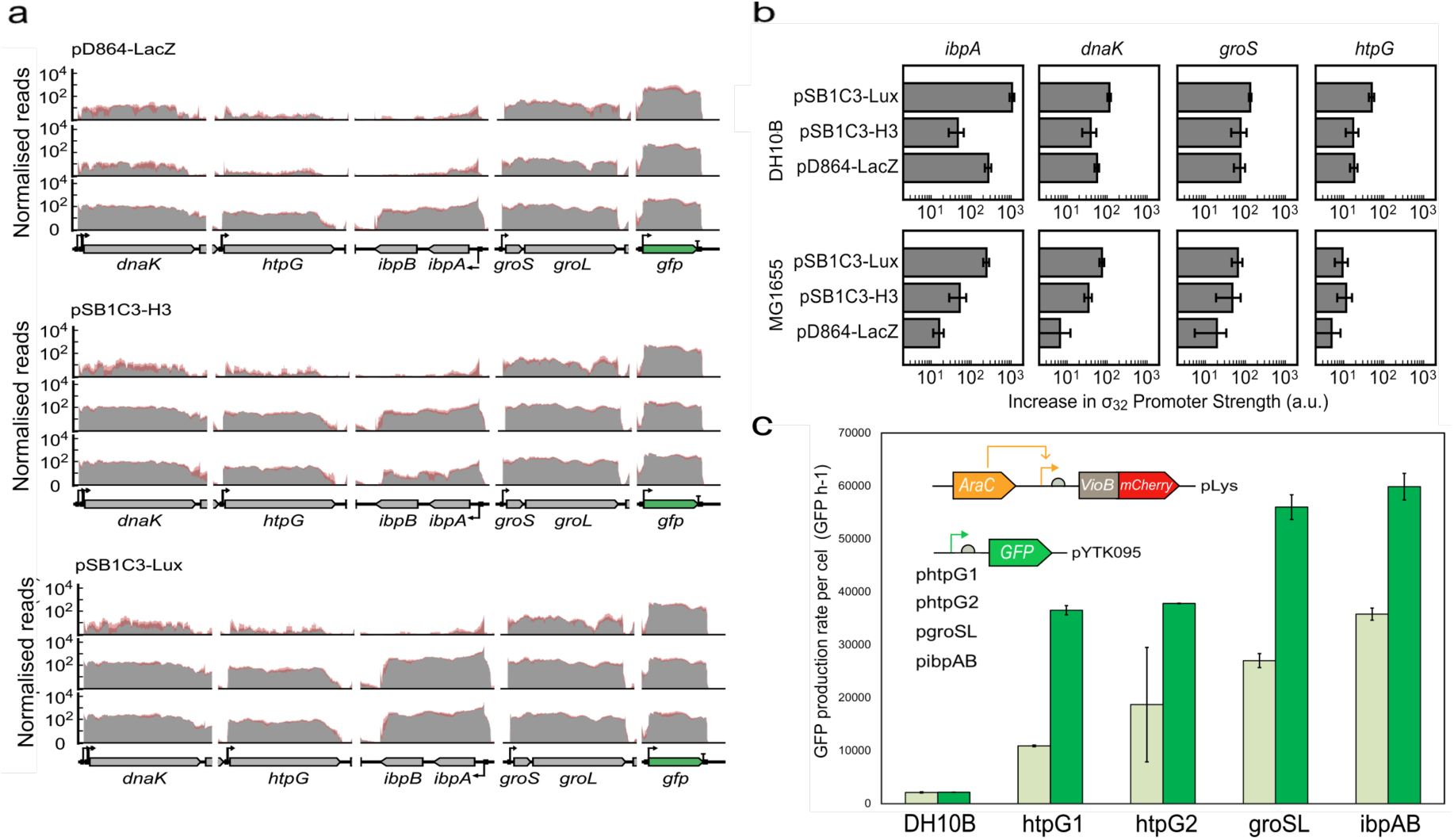
Characterisation of burden-dependent activation of genomic and plasmid-based σ32-regulated promoters. **(a)** Transcription profiles of the *dnaKJ*, *htpG*, *ibpAB and groSL* regulons and the GFP monitor cassette from DH10B cells with induced synthetic constructs pD864-LacZ, pSB1C3-H3 and pSB1C3-Lux. Traces show normalised reads mapping to the plasmid DNA. Top profiles are for strains containing only the empty plasmid 15 min post-induction, middle and bottom profiles are for strains containing synthetic constructs 15 and 60 min post-induction, respectively. Solid grey regions show average and transparent red regions span the min to max of transcription profiles generated from three biological replicates. Genetic designs below each profile are drawn using SBOLv notation and show the precise location of each gene and key promoters in the genome. **(b)** Transcriptional response of genomic *ibpAB*, *dnaKJ*, *groSL* and *htpG* promoters. The increase in transcription strength in response to construct expression from the genomic *ibpAB*, *dnaKJ*, *groSL* and *htpG* promoters in DH10B and MG1655 strains after 60 minutes as calculated from the RNAseq data. Error bars show standard deviation of three independent repeats. **(c)** The *in vivo* response to burden from the *htpG1, htpG2, groSL* and *ibpAB* promoters. All four promoters were assessed for their ability to induce GFP expression from a high-copy plasmid when co-transformed with a construct expressing VioB-mCherry upon arabinose induction. Data show the GFP production rate per cell from the four promoters 2 hour post-induction of VioB-mCherry expression in DH10B cells. Dark and light green bars represent uninduced (no arabinose) and induced samples (arabinose), respectively. Error bars show standard deviation of two independent repeats.

With a view to utilising these native promoters we next assessed whether their burden-based induction can be recreated when taken out of their natural genomic context and used to express GFP from a plasmid. For the *groSL* and *ibpAB* promoters this simply required straightforward cloning, however the *htpG* and *dnaK* promoters naturally consist of two and three overlapping promoters, respectively, making their cloning more complicated. For *htpG*, it was possible to separate the two overlapping promoters (*htpG1* and *htpG2*) and test these (**Table S5**). However, the *dnaK* promoter could not be similarly separated into functional promoters and so was not explored further (see **Supplementary Note 1**). The four cloned promoters were then assessed for their ability to drive GFP expression in DH10B cells in the presence of a burden-causing VioB-mCherry expression construct when it is uninduced and induced (**Figure 3c**, also see **Figures S7** and **S8**). All four promoters showed basal GFP expression when the VioB-mCherry construct was uninduced, but significantly increased expression after its induction, confirming that their response to burden could be achieved when taken out of their genomic context.

While the *ibpA* and *groS* promoters showed the greatest expression in response to burden, the *htpG1* promoter displayed the best on/off characteristics and so we considered it to have the greatest utility as a broad, indirect sensor for the burden of expressing synthetic constructs in *E. coli*. RNAseq analysis showed activation of this promoter in both strains for all constructs despite there being substantial differences in their genetic content, their inducers, what proteins they produce and their plasmid backbones. As well as being a potential biosensor for burden, the *htpG1* promoter offers a key component for creating a negative feedback system that controls gene expression burden in *E. coli*: burden is sensed through the *htpG1* promoter, and in response an effector molecule is produced that represses the expression of the genes that cause this burden. A feedback system with such design should enable robust control of host capacity, regardless of growth conditions and construct design (e.g. promoter strength, RBS strength, protein coding sequence).

For this system to be as general as possible, we generated a negative biomolecular feedback controller construct where the *htpG1* promoter drives CRISPR guide RNA (sgRNA) expression, which, in turn, directs the binding of dCas9 to target sequences within specific promoter regions in order to sequence-specifically repress their expression. For this to have as rapid a response as possible the construct was designed to express dCas9 constitutively (**Table S6**), so that repression can occur as soon as the sgRNA is transcribed from the burden-induced *htpG1* promoter (**Figure 4a**). While constitutive dCas9 expression represents an extra cost to the host cell, we found that this was minimal when expressed appropriately at a low enough level (**Figure S9**). Thus, the negative feedback was encoded onto a medium copy plasmid and its design conceived to be modular and so easily adapted to any synthetic construct. Modularity of the system allows easy replacement of the *htpG1* promoter or the sgRNA via specific rare restriction sites flanking these parts (**Figure S10**).

**Figure 4.**
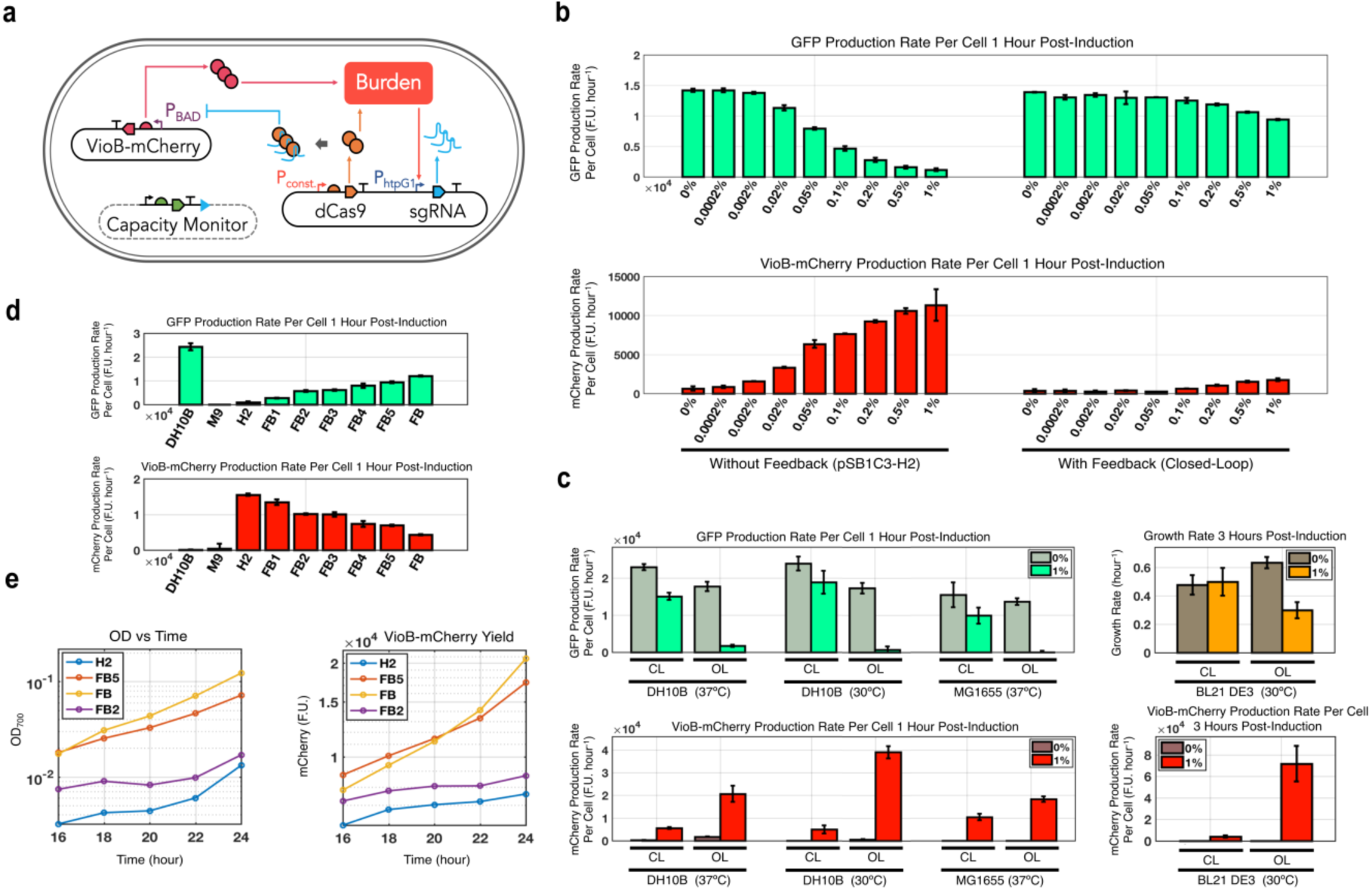
Burden-driven feedback control system. **(a)** Schematic of the feedback system. The dCas9-sgRNA feedback controller was placed on a medium copy plasmid and co-transformed in DH10B monitor cells with a construct (pSB1C3-H2) producing VioB-mCherry at high levels from a pBAD promoter. In the feedback construct the htpG1 promoter is placed upstream of a sgRNA targeting the pBAD promoter. On the same plasmid a constitutively-expressed dCas9 cassette is encoded. When VioB-mCherry is produced and burden is consequently triggered in the cell, the *htpG1* promoter activates production of sgRNA. The sgRNA-dCas9 complex binds to and represses pBAD decreasing VioB-mCherry production. **(b)** Functionality of the feedback. GFP (top) and VioB-mCherry (bottom) production rate per cell are shown when different levels of arabinose are used to trigger expression in DH10B. Error bars show standard deviation of three independent repeats. **(c)** Robustness and portability of the feedback. The biomolecular feedback was tested in DH10B at 37°C and 30°C and MG1655 cells at 37°C with integrated capacity monitor (left panels) and in BL21-DE3 cells at 37°C (right panels). GFP (top) and VioB-mCherry (bottom) production rate per cell are shown for DH10B and MG1655 when 0% and 1% of arabinose are added. For BL21-DE3, growth rate (top) and VioB-mCherry rate (bottom) are shown instead. Error bars show standard deviation of three independent repeats. **(d)** Tunability of the feedback system. A library of feedback controllers was designed where single point mutations were inserted in the sgRNA sequence in order to decrease its binding affinity to the pBAD promoter. GFP (top) and VioB-mCherry (bottom) production rate per cell 1 h post induction for the library are shown. Error bars show standard deviation of three independent repeats. **(e)** 24 hour time-course experiment. Constructs FB, FB2 and FB5 from (d) were chosen to run a 24 hour batch growth time course after addition of 1% arabinose to the cultures containing the feedback system and pSB1C3-H2 construct. The culture density (left) and total VioB-mCherry yield (right) in growth flasks are compared from 16 to 24 hours.

The feedback controller was first tested in DH10B cells with the genomically-integrated GFP capacity monitor in the presence of construct pSB1C3-H2, a previously designed construct similar to pSB1C3-H3 with a stronger RBS and so even higher burden ^16^. The *htpG1* feedback controller was designed so that the expressed sgRNA targets dCas9 to the core region of the pBAD promoter to inhibit transcription of VioB-mCherry. In *E. coli* without the feedback controller (pSB1C3-H2 construct), as the level of arabinose inducer in the media is increased, the rate of VioB-mCherry production per cell rises and the capacity for host expression (as measured by GFP production rate per cell) decreases. With the feedback plasmid present, the VioB-mCherry production is kept low even at the highest arabinose concentrations, while the host capacity remains high (**Figure 4b**). This demonstrates that the feedback controller can effectively limit the output (VioB-mCherry) over a range of induction levels in order to maintain high host expression capacity.

To test the robustness of the controller, we verified that its functionality would persist under different environmental conditions by growing the cells at different temperatures (**Figure 4d and Figure S11**). To also prove the general portability of the system we moved it in different genetic backgrounds demonstrating its continued robust performance in both MG1655 and BL21-DE3 *E. coli* strains (**Figure 4d, Figure S12, Figure S13**).

As applications of this feedback may not always require having high host capacity at the cost of low construct expression, we next looked at whether the strength of feedback could be tuned. Indeed, different applications may require more construct output in exchange for sacrificing host capacity. To tune the trade-off between construct production and host capacity, we designed mismatches into the targeting sequence of the sgRNA in the feedback construct (**Table S6**). Mismatches reduce the affinity of the dCas9-sgRNA complex for its target promoter and thus decrease the strength of repression ^24^. Random mutations at single positions within the sgRNA targeting sequence allowed to modify the feedback gain, thereby producing a series of feedback controllers that give increasing pSB1C3-H2 output at the cost of decreasing host capacity (**Figure 4c**). Thus, this simple approach of adding mismatches into the targeting sequence enables the creation of a library of feedback controllers that offer different maximum construct outputs and resulting host cell capacities.

DH10B cells containing the pSB1C3-H2 construct along with two of the mismatched feedback controllers (FB2 and FB5) or with the original fully-repressing feedback controller (FB) were then assessed in batch cultures over 24 hours and compared against DH10B cells with the pSB1C3-H2 construct alone (**Figure 4e**). As a proxy for the total production of the synthetic gene product (VioB-mCherry) from all four samples, the total red fluorescence of the culture was measured. Up to 16 hour of growth, the most productive culture in terms of total VioB-mCherry product per flask was the strain with mismatch feedback controller FB5. However, after 24 hours, the culture with the strongest feedback gain (FB) yielded the largest amount of total product, presumably as this strain grew the fastest over the 24 hour period and, therefore, had more total biomass per flask. Thus, while the strains with these feedback controllers exhibit lower VioB-mCherry production rates per cell, they result in larger total production over time as their increased capacity for expression enables significantly improved growth of the culture.

## DISCUSSION

By combining our *in vivo* assay with multiplex RNAseq analysis we here revealed a host response to expression burden while determining the transcriptional impact of synthetic construct expression. This was done by investigating the burden imposed on two *E. coli* strains when expressing three exemplar inducible constructs that have different DNA sequences and functions. Regardless of their genetic content, all constructs consumed a significant fraction of the cell’s transcriptional resources, reduced host gene expression capacity and triggered upregulation of σ32-regulated promoters within the first hour. We exploited this latter observation to build a general and tunable biomolecular feedback controller system with modular design that can respond to burden and use dCas9-mediated repression to maintain host expression capacity to desirable levels. We showed this system to be portable, robust to different conditions and able to improve protein production yield in 24 hour batch experiments. We anticipate wide use of this feedback system in future work.

The use of RNAseq in particular provided new insights into how cells respond to sudden overexpression of genes in non-native contexts, and the data obtained here provides new information towards understanding cellular resource reallocation and how the burden of extra expression eventually leads to reduced bacterial growth rates. It also highlighted how significant the impact transcription of plasmid constructs is on host cells. One hour post-induction, between 20% and 50% of all mRNAs in our RNAseq samples mapped to the genes from the synthetic constructs. Given that mRNA production in growing *E. coli* is estimated to represent approximately half of all active transcription (the other half being rRNA and tRNA production) ^25^ this means that constructs with very high burden could be using up as much as a quarter of all transcriptional resources, such as RNA polymerases, when expressed. Understandably this reallocation of resources will also have a major impact on translational resources, especially if the very high numbers of mRNAs expressed from synthetic constructs have strong RBS sequences that rapidly recruit ribosomes and have long, difficult-to-translate sequences that will sequester ribosomes away from the pool required for host gene expression ^14^. Future work to assess the impact of burden on translational resources could extend from our transcriptome analysis by making use of RiboSeq, an adaptation to RNAseq protocols that quantifiably determines the footprints of all ribosomes on mRNAs in cells ^26^.

Similarly, other recent RNAseq advances could be used to extend our work. The use of spike-in sequences ^27^ offers a route towards converting RNAseq data into true transcript numbers per cells. And the recently described TagSeq method ^28^ for multiplex RNAseq now greatly simplifies the laborious sample prep of multi-sample RNAseq and will accelerate future attempts to repeat our approach to analyse many other different synthetic constructs. Indeed, perhaps the greatest limitation of the work done here is that the sequencing of 90 samples only results in the analysis of three different types of synthetic constructs, albeit with high confidence in results and controls (n=3) and with two time-points in two strains. For a more complete picture on the transcriptome response to burden it would be desirable downstream to assess many other synthetic constructs, including those with dynamic or switchable expression states (e.g. oscillators, logic gates), those present at lower copy-numbers (e.g. on low copy plasmids or chromosomally-integrated) and those encoding more diverse functions (e.g. secreted proteins, membrane proteins, RNA-only devices).

However, despite the small set of synthetic constructs we assessed here we covered enough diversity to identify a general burden response. The results for pD864-LacZ are particularly interesting, as this construct uses a different inducer, plasmid backbone and antibiotic selection to the other constructs and does not express any foreign genes or immediately express proteins at overwhelming levels. However, even with this construct σ32-responsive promoters are triggered within an hour of induction in both *E. coli* strains. Thus the heat-shock stress response, known to be associated with overexpression of recombinant proteins ^29^, is also seen when native proteins are expressed in non-native contexts. Based on this observation we would expect these same promoters to activate in response to burdensome gene expression of almost any protein in *E. coli*, and that codon-optimisation would not solve this problem (native *lacZ* is presumably already codon-optimised). The σ32 response is clearly a rapid and sensitive mechanism for the cells to adapt resources in the face of unexpected gene expression ^30 31^. It triggers chaperone production to prevent protein misfolding and aggregation as well as proteases to degrade unfolded peptides and rescue stalled ribosomes. This response therefore matches expectations that expression burden is most significant at the stage of protein translation.

Of the native σ32-responsive promoters we analysed, the *htpG1* promoter provided the best on/off characteristics when expressed in a plasmid context and as such is the basis for our biomolecular feedback controller. The same promoter could also be used in other burden-triggered systems, for example by activating fluorescent reporter proteins or selectable markers in order to rapidly screen construct libraries for or against burdensome expression. Our chosen design for the feedback controller was intentionally as modular and general as possible, and can instead incorporate different response promoters, for example to generate controllers that respond to other stresses (e.g. lack of metabolites) or other phenotypes that could be investigated as done here with RNAseq.

Specifically, the use of CRISPR/dCas9 allows the system to theoretically regulate any gene target simply through expression of guide RNAs that match the target core promoter. Future work could examine burden-based regulation of a wider diversity of synthetic constructs with different promoters and proteins being expressed. Multiple guide RNAs could also be expressed in parallel if more than one target requires regulation and the system could be switched to upregulate expression rather than repress by fusing a transcriptional activation domain to the dCas9 protein. Importantly, because the only burden-activated component is the guide RNA, the system dynamics are quick as no new translation is required to produce the regulator. This speed plus the ease of retargeting and tuning, means that this system is a major improvement on existing gene expression feedback control devices that rely on the slow and costly translation of repressor proteins that can only target their cognate promoters ^32^. A more complete understanding of feedback dynamics could be achieved by assessing the ability of our system to adjust expression during fluctuating conditions (e.g. in chemostat experiments). Furthermore, future iterations of our system could place the feedback controller (or just the dCas9 gene) into the strain genome to further lower its impact on the host cells. However, in its current plasmid-based form it provides a simple extra construct that can be added alongside synthetic constructs into a user’s strain of interest without these strains needing modification.

## METHODS

### Bacterial strains

Strains MG1655 (K-12 F-λ*-rph*-1), DH10B (K-12 F- λ*- araD139* Δ*(araA-leu)7697* Δ*(lac)X74 galE15 galK16 galU hsdR2 relA rpsL150*(StrR) *spoT1 deoR* φ*80*d*lacZ*ΔM15 *endA1 nupG recA1 e14-mcrA* Δ*(mrr hsdRMS mcrBC)*) were obtained from the National BioResource Project (NBRP) Japan. Strain BL21(DE3) 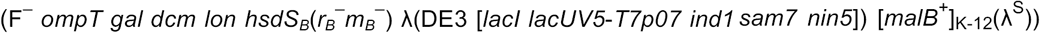 was obtained from NEB. The capacity monitor cassette consisted of a synthetic strong constitutive promoter, a synthetic RBS, a codon-optimized superfolder GFP coding sequence and a synthetic unnatural bi-directional terminator. Details of construction and validation of the capacity monitor used in this paper were previously published ^16^.

### DNA constructs design

The four expression constructs giving inducible heterologous gene expression are illustrated in **Figure 1b**. Construction of constructs pSB1C3-H3, pLys-M1 and pSB1C3-Lux was described previously ^16^. Construct pD864-LacZ was obtained by PCR amplification of the LacZ coding sequence from *E. coli* genome and restriction digestion into the pD864-SR plasmid purchased from DNA2.0. Control plasmids were the high-copy, chloramphenicol-selectable pSB1C3 plasmid obtained by restriction digest and religation; the medium-copy, chloramphenicol-selectable pLys plasmid and the high-copy, ampicillin-selectable pD864 plasmid, both obtained by PCR amplification and religation to remove the synthetic construct insert.

### Construction of plasmid based htpG1, htpG2, groSL and ibpAB promoters

Restriction digestion and religation were used after PCR amplification of plasmid YTK095 containing a GFP coding sequence, Ampicillin resistance and pMB1 origin of replication. PCR was used to add SfiI and PacI restriction sites at the 5’ and 3’ of the plasmid, respectively. gBlocks for each of the promoters were ordered from IDT containing the SfiI and PacI restriction sites upstream and downstream of the promoter sequence, respectively (**Table S5**).

### Construction of the burden-based feedback system

The sequence of the dCas9 protein was PCR amplified from a commercial plasmid and inserted on the medium copy plasmid pZA16 carrying ampicillin resistance downstream of the constitutive BBa_J23113 promoter. The reverse primer for dCas9 amplification was designed so that it could bear SfiI/PacI/AscI restriction sites. A GeneString containing the htpG1 promoter sequence was ordered from GeneArt with the SfiI and PacI restriction sites at the 5’ and 3’ respectively. The sgRNA targeting pBAD promoter was ordered from GeneArt with PacI and AscI restriction sites at the 5’ and 3’ respectively. Both the promoter and the sgRNA were inserted downstream of dCas9 using restriction digestion and riligation. All enzymes used for cloning were obtained from NEB.

Inverse PCR was used to create a library of randomly mutated promoters starting from BBa_J23113 upstream of dCas9. The sequence of the promoter selected for final implementation of the feedback system is shown in **Table S6**.

Construction of the library of randomly point-mutated sgRNA was done using inverse PCR with insertion-encoding 5’ phosphorylated primers, followed by DpnI digestion and religation before transformation in DH10B cells. Six PCR reactions were carried out separately with six forward primers each carrying a mutation at a different position. All PCRs were carried out with the NEB Phusion High Fidelity Polymerase. Co-transformation of the library with H2 was then performed in DH10B. Seventeen colonies were picked and sent to DNA sequencing for identification of the position of the point-mutation in the sgRNA of each of these colonies. The constructs were then co-transformed together with pSB1C3-H2 in DH10B cells. Sequences of the sgRNA library members are shown in **Table S6**.

Construction of the open-loop counterpart of the feedback plasmid was carried using inverse PCR to add BsaI sites around the htpG1 promoter and the sgRNA targeting pBAD. A 272-bp gBlock was ordered and synthesized by IDT. The gBlock contains BsaI sites around the htpG1 promoter and the random sgRNA (**Table S6**) was designed using R2o DNA Designer ^33^. The two strings were assembled via Golden Gate Assembly, and the resulting plasmid was transformed in DH10B monitor cells. The construct was verified by DNA sequencing before use. The open-loop version was then co-transformed together with pSB1C3-H2 in DH10B monitor cells.

### Burden assay and RNAseq time course

For the burden assay and time course *E. coli* cells with construct and control plasmids were grown at 37°C overnight with aeration in a shaking incubator in 5 ml of defined supplemented M9 + Fructose media with the appropriate antibiotic. In the morning, 60 μl of each sample was diluted into 3 ml of fresh medium and grown at 37°C with shaking for another hour (outgrowth). 200 μl of each sample were then transferred in 8 wells of a 96-well plate (Costar) at approximately 0.1 OD (600 nm). The samples were placed in a Synergy HT Microplate Reader (BioTek) and incubated at 37°C with orbital shaking at 1,000 rpm for 1 h, performing measurements of GFP (excitation (ex.), 485 nm; emission (em.), 528 nm) and OD (600 nm) every 15 minutes. 60 minutes into the incubation, the plate was briefly removed so inducer could be added to wells, and this time point was set as time 0. In the burden assay, cells were let to grow in the reader for 4.5 hours with measurements of GFP [excitation (ex.), 485 nm; emission (em.), 528 nm] and OD (600 nm) every 15 minutes. In the RNAseq analysis, samples were instead removed from wells at 15 and 60 minutes after induction for processing. Specifically, 170 μl were taken from each of four wells per time point and collected into a fresh tube were 1.360 ml of RNA protection buffer had previously been added. Samples were left for 5 minutes at RT and then centrifuged at 4°C at maximum speed. Supernatant was discarded and pellets frozen at -20°C. Three replicates were repeated independently on three different days for a total of ninety samples used to produce the final data set (7 constructs X 2 strains X 3 replicates X 2 time points = 84 samples; plus DH10B:GFP cells X 3 replicates X 2 time points) (**Table S2**). To avoid inhibition of arabinose-induced expression due to catabolite repression, 0.4% fructose was used as the main carbon source. Inducers were added to appropriate wells at 0 minutes. Final inducer and antibiotic concentrations used in assays were as follows: L-arabinose, 0.2%; L-Rhamnose, 2%; ampicillin, 100 μg/ml; and chloramphenicol, 34 μg/ml.

### Data analysis

For the plate-reader characterisation assays, the growth and the protein expression rates per hour were calculated following the procedure of ^16^. Growth rate at t2 = [ln(OD(t3))– ln(OD(t1))]/(t3–t1), GFP capacity rate at t2 = [(total GFP(t3) – total GFP(t1))/(t3–t1)]/OD(t2), and mCherry output rate at t2 = [(total mCherry (t3)–total mCherry (t1))/(t3–t1)]/OD(t2), where t1 = time-15 min, t2 = present time, and t3 = time+15 min. Mean rates and their standard errors were determined from the three biological replicates from the same 96-well plate. 400 was added to all mCherry output rates to account for the background red fluorescence of M9 + fructose media, which decreases at a rate of approximately 400 h^-1^ as the media is consumed by cells during growth ^16^.

### RNAseq library preparation

RNA extraction was performed using RNeasy Mini Kit from Qiagen [Cat No 74104]. To remove possible traces of genomic DNA contamination, 2 μg of each sample were treated for a second time with DNAseI from Qiagen [Cat No 79254]. Total RNA quality and integrity was assessed using Agilent 2100 Bioanalyzer and Agilent RNA 6000 Nano kit [Cat No 5067-1511]. Samples had an average RIN of 9.5. Enrichment of mRNA was performed using MicrobExpress rRNA removal kit from Thermo Scientific [Cat No AM1905] ^22^. Successful rRNA depletion was assessed with analysis on Bioanalyzer. Retrotranscription was then performed starting from 50 ng total enriched mRNA using Tetro cDNA synthesis kit from Bioline [Cat No BIO-65043] and 6 μl of Random Hexamers [Cat No BIO-38028] per reaction. Second cDNA synthesis was performed adding to the first strand synthesis mix 5 μl of Second strand synthesis buffer [Cat No B6117S], 3 μl of dNTPs [Cat No N0446S], 2μl of RNAseH [Cat No M0297L] all from NEB, 2 μl of Polymerase I from Thermo Scientific [Cat No 18010025] and 18 μl of water, per reaction. Samples were incubated at 16°C for 2.5 h. Purification of cDNA was performed using MiniElute PCR purification kit [Cat No 28004] with final elution in 10 μl of DEPC-treated free water. cDNA was quantified using a Qubit fluorometer (Invitrogen). Library preparation was performed using the Nextera XT kit from Illumina [Cat No FC-131-1096] starting from 1 ng of total cDNA. The original protocol was modified where 3 min tagmentation and 13 cycles of step-limited PCR were used. Ampure beads from Beckman Coulter [Cat No A63880] were used for library purification. Library quality assessment and quantification was performed with Agilent 2100 Bioanalyzer and Agilent high sensitivity DNA analysis kit [Cat No 5067-4626]. Finally all 90 samples were pooled together in the same reaction tube at a final concentration of 1 nM.

### RNAseq library sequencing

Library sequencing was performed at the Imperial College London Genomic Facility. Two lanes from the HiSeq 2500 Sequencer were used for paired end sequencing with read length of 100 bp.

### Sequencing quality control and alignments

Raw reads for all sequenced samples were quality assessed and trimmed using Trim Galore v0.4.1 with default settings. After assessment for potential batch effects the technical replicates were pooled. *E. coli* strain (DH10B and MG1655) sequences were obtained from Ensembl genomes release 31. A FASTA format sequence file corresponding to the composite of strain, plasmid and integrated GFP was constructed for each sample, and used as a reference for read alignment. Trimmed reads were aligned using BWA mem algorithm v0.7.12-r1039 using default settings. Samtools v1.3.1 was used on resultant alignments to create a sorted BAM file for each sample. The biological replicates were checked for any batch effects before generating the raw counts using Bioconductor Rsubread package v1.12.6. The normalised FPKM counts were generated using Bioconductor edgeR package version 3.4.2 accounting for gene length and library size (by applying TMM normalisation), which were used for downstream analysis.

### Transcription profiles and promoter characterisation

The method described in Gorochowski *et al.* ^34^ was used to generate the transcription profiles from RNAseq data. Raw reads from the sequencer in a FASTQ format were mapped to the appropriate *E. coli* host genome reference sequence (NCBI RefSeq: NC_000913 for MG1655, and NCBI RefSeq: NC_010473.1 for DH10B) with separate reference sequences included for the GFP monitor and appropriate plasmid constructs using BWA version 0.7.4 with default settings. These BAM files were then separately processed by custom Python scripts to extract the position of the mapped reads, count read depths across the reference sequences, and perform corrections to the profiles at the ends of transcription units. These profiles were then normalised to enable comparisons between samples. Characterization of promoters was performed using custom Python scripts as in Gorochowski *et al.* ^34^, which took as input a GFF reference of the construct defining the location of all parts. Visualisations of the transcription profiles and associated genetic design information was generated in an SBOL Visual ^35^ format using DNAplotlib version 1.0 ^36^. All analyses were carried out using custom scripts run using Python version 2.7.12, NumPy version 1.11.2, and matplotlib version 1.5.3.

### Differential gene expression

DESeq2 was used for differential expression analyses ^37^. Gene expression was compared between cells transformed with synthetic constructs and the analogous cells transformed with the corresponding empty plasmid. Reads mapping to ribosomal genes or to the synthetic constructs were excluded. Differentially expressed genes were annotated with data extracted from the EcoCyc database ^38^ using custom Python code.

### Testing the robustness of the feedback system

For plate-reader assays to test the robustness of the feedback system to arabinose-induction perturbation, the constructs were induced with final concentrations of arabinose: 0%, 0.0002%, 0.002%, 0.02%, 0.05%, 0.1%, 0.2%, 0.5%, and 1%.

### Shake-flask–scale growth

Constructs pSB1C3-H2, FB+H2, FB2+H2 and FB5+H2 were assessed in DH10B over 24 hours of exponential growth in M9 fructose (0.4%) media with 1% final concentration of arabinose. The experiment was done as previously described ^16^. Starter cultures of pSB1C3-H2, FB+H2, FB2+H2 and FB5+H2 were taken on plate from individual colonies, and used to inoculate 3 ml of M9 fructose media, supplemented with the appropriate antibiotics, in 15 ml culture tubes. The cultures were then grown in the 37°C shaking incubator for 5 hours, before being diluted to 0.015 OD600. 50 μl of the diluted culture (∼150,000 cells) was used to inoculate batch cultures of 50 ml supplemented M9 with 0.4% fructose, 1% L-arabinose, and the appropriate antibiotics: ampicillin, 100 g/ml (50 g/ml when both ampicillin and chloramphenicol are supplementing the medium); and chloramphenicol, 34 g/ml (17 g/ml when both ampicillin and chloramphenicol are supplementing the medium) in 500-ml baffled shake flasks. The cultures were then grown in the 37°C shaking incubator during 16 h, at which point 200 μl of each culture was dispensed in individual wells of a 96-well plate, and 350 μl was diluted in 650 μl of PBS and stored in the fridge every hour from 16 hours until 24 hours. The 96-well plate was placed in a preheated plate reader at 37°C every hour to perform OD measurements (OD600 and OD700), GFP measurements (485 nm (excitation)/528 nm (emission)) and mCherry measurements (590 nm/645 nm) every 2 min. Only the first five measurements were taken and averaged to obtain the OD, GFP and mCherry values at the specific time point.

## AUTHOR CONTRIBUTIONS

FC, GBS and TE designed the research; FC, AB, CG, AW performed the experiments; FC, SF, TG, YL, GBS and TE analysed the data; FC, SF, TG, OB, GBS and TE wrote the paper.

## ACKNOWLEDGEMENTS

FC was supported by EPSRC grant EP/J021849/1. TEG was supported by BrisSynBio, a BBSRC/EPSRC Synthetic Biology Research Centre (grant BB/L01386X/1). AB, OB and CG were supported by EPSRC grant EP/M002306/1. YNL was supported by the NIHR Imperial Biomedical Research Centre. AW was supported by BBSRC grant BB/K006290/1. GBS was supported by EPSRC Fellowship EP/M002187/1, and grants EP/J021849/1 and EP/P009352/1. TE was supported by EPSRC Fellowship EP/M002306/1 and grant EP/J021849/1.

